# Intracellular bio-synthesizing Gold Nanoparticles using bacterial systems and their applications in photothermal therapy (A green approach)

**DOI:** 10.64898/2026.11.07.765052

**Authors:** Gul Naz Ashraz, Marta Palau Gauthier, Qiutian She, Pilar Rivera-Gil, Javier Macía

## Abstract

**Background:** Conventional cancer therapies are heavily constrained by systemic toxicity, off-target side effects, and poor deep-tissue penetration due to the highly heterogeneous tumor microenvironment (TME). Although gold nanoparticles (AuNPs) possess exceptional localized surface plasmon resonance (LSPR) properties ideal for photothermal therapy (PTT), their clinical translation is bottlenecked by inconsistent passive accumulation and chemical toxicity arising from conventional synthetic surfactants.

**Objective:** This study developed a sustainable, biocompatible bio-hybrid platform utilizing the probiotic *Escherichia coli* Nissle 1917 (EcN) as an autonomous, tumor-homing living carrier integrated with AuNPs synthesized via an eco-friendly intracellular green chemistry approach.

**Methods:** EcN cultures were prepared in Luria-Bertani (LB) medium and subsequently exposed to a 1 mM chloroauric acid (HAuCl_4_) metallic precursor. A facile, room-temperature green synthesis protocol was established by utilizing ascorbic acid (Vitamin C) as a non-toxic co-reductant to drive a seed-mediated crystallization onto intra-bacterial metallic seeds. The bio-hybrid system (EcN::AuNPs) was characterized via visual inspection, UV-Vis spectroscopy, and *in vitro* PTT assays under continuous-wave near-infrared (NIR) laser irradiation.

**Results:** Visual confirmation showed an irreversible color shift from pale yellow to a deep dark-brownish hue, serving as a direct diagnostic indicator of successful Au^3+^→Au^0^ reduction. UV-Vis spectroscopic analysis revealed a distinct, broadened surface plasmon resonance (SPR) band centered at 560nm, indicating the formation of stabilized, isotropic gold nanostructures (30-60nm) subjected to near-field electromagnetic coupling within the complex intracellular compartments (periplasm and cytoplasm). Biocompatibility assays demonstrated a significant collapse in cell viability from 10^8^CFU/mL down to 10^3^CFU/mL, driven by a combination of trivalent gold heavy-metal toxicity, mechanical disruption of the peptidoglycan wall from spatial confinement, and severe metabolic/oxidative stress. Nevertheless, the remaining surviving population maintained required active chemotactic capabilities, while the lysed bacterial matrix provided a natural protein capping layer that prevented nanoparticle aggregation. Under 5 minutes of continuous NIR laser irradiation, the EcN::AuNPs platform exhibited exceptional concentration-dependent hyperthermia kinetics, triggering a drastic thermal escalation from 20°C to an absolute peak of 75.4°C (ΔT=55.4°C), vastly outperforming all baseline control groups (pure EcN, bare HAuCl_4_, and standalone ascorbic acid).

**Conclusion:** The engineered EcN::AuNPs bio-hybrid platform represents a highly efficient and biosafe “bacteriobot” paradigm. By shifting the SPR footprint and leveraging the native bacterial cell envelope for colloidal stability, this system achieves an optimal equilibrium between active biological tumor targeting and rapid, localized photothermal ablation, offering a promising alternative for targeted oncological interventions.

## Introduction

Cancer remains one of the leading causes of mortality worldwide, accounting for a significant percentage of global deaths annually[97], [98], [99]. Despite extensive biomedical advancements, the clinical management of cancer is severely challenged by its highly heterogeneous nature, which prevents a “one-size-fits-all” therapeutic approach. Conventional treatment modalities, primarily chemotherapy, radiotherapy, surgery, and immunotherapy, frequently suffer from systemic limitations. These include severe off-target toxicity to healthy tissues, profound suppression of the immune system, and the critical risk of inducing secondary malignancies[100], [101], [102], [103], [104], [105]. Consequently, there is an urgent and ongoing imperative to develop targeted, highly efficacious, and minimally invasive therapeutic alternatives[106].

In this context, gold nanoparticles (AuNPs) have emerged as a premier platform in nanomedicine [107], [108]. Characterized by dimensions typically ranging from 1 to 100 nm, AuNPs exhibit unique optical, electronic, and physicochemical properties that diverge drastically from their bulk counterparts due to quantum confinement and surface effects[109]. Among metallic nanoparticles, AuNPs are exceptionally prominent owing to their remarkable biocompatibility, high chemical stability, and versatile surface chemistry, which allows for precise functionalization[110], [111]. These tailored attributes have facilitated their widespread application in biomedical engineering, spanning drug delivery vehicles, biosensing, molecular diagnostics, and direct oncological therapeutics[112], [113], [114]. Notably, AuNPs are uniquely suited for photothermal therapy (PTT) due to their ability to exploit the enhanced permeability and retention (EPR) effect for passive tumor accumulation, as well as active targeting via surface ligands, followed by activation using near-infrared (NIR) laser light[115], [116].

The therapeutic rationale behind PTT relies on the differential thermal sensitivity between malignant and healthy tissues; cancer cells are notoriously susceptible to hyperthermia due to the hypoxic, acidic, and nutrient-deficient microenvironment characteristic of solid tumors [117]. When engineered into specific morphologies, such as nanospheres, nanoshells, nanorods, or nanostars, AuNPs display strong localized surface plasmon resonance (LSPR)[118], [119]. Literature demonstrates that upon localized accumulation within the tumor site, irradiation with a light source tuned to the AuNPs’ LSPR peak triggers a rapid and efficient conversion of absorbed photon energy into localized thermal energy[120][121]. The efficacy of this photothermal conversion is dictated by the specific absorption cross-section of the nanoparticles and their light-to-heat conversion efficiency. Typically, NIR lasers (700–1350 nm) are preferred as excitation sources because biological tissues exhibit minimal absorption in this “optical window,” enabling deep tissue penetration with negligible collateral damage[122], [123]. The foundational efficacy of AuNPs in mediating thermally induced cellular death was pioneeringly demonstrated by Pitsillides et al. (2003), who utilized anti-CD8-labeled gold nanoparticles to selectively target and destroy specific T-cell populations[124], paving the way for modern targeted nanoplasmonic hyperthermia.

### 3. Advanced Targeting Strategies: Probiotic-Mediated Delivery of AuNPs

Despite the exceptional photothermal properties of gold nanoparticles (AuNPs), their clinical translation is heavily bottlenecked by the challenge of precise targeted delivery. Passive targeting relies primarily on the enhanced permeability and retention (EPR) effect, which frequently suffers from inconsistent tumor accumulation, high splenic and hepatic clearance, and off-target toxicity in healthy tissues[125], [126] [127], [128]. Consequently, contemporary research has pivoted toward designing “smart” active delivery platforms capable of autonomously traversing complex biological barriers and selectively homing into neoplastic lesions. Among the most disruptive paradigms in this domain is the utilization of live engineered probiotics as autonomous cellular carriers, an approach often termed bacterial-mediated tumor targeting or “bacteriobots”[129], [130], [131], [132].

Traditional non-living nanocarriers lack the ability to actively navigate the dense extracellular matrix of solid tumors, resulting in poor deep-tissue penetration [133], [134], [135]. In contrast, living drug delivery systems can actively target, penetrate, and colonize tumor sites. Certain non-pathogenic bacterial strains, most notably *Escherichia coli* Nissle 1917 (EcN), exhibit a natural, intrinsic tropism for solid malignancies. When administered systemically or orally, these micro-organisms preferentially colonize and proliferate within the tumor microenvironment (TME) [136], [137], [138]. This selective colonization is driven by the unique physicochemical hallmarks of the TME, such as chronic hypoxia, chaotic angiogenesis, extensive necrosis, and immune-suppressed niches, which provide an ideal anaerobic or microaerophilic sanctuary for bacterial growth while shielding them from immune clearance[136], [137], [139].

Beyond their innate homing and motility capabilities, probiotics like EcN serve as versatile synthetic biology chassis. They can be genetically engineered to express therapeutic macromolecules, cytokines, or tumor-specific antigens locally, or physically/metabolically modified to act as microscopic “Trojan horses” that transport inorganic payloads, such as AuNPs, directly into the necrotic core of the malignancy. This bio-hybrid delivery strategy offers unprecedented precision for gastrointestinal oncology, particularly colorectal cancer (CRC). In CRC, the oral or local administration of AuNP-loaded probiotics bypasses many of the systemic vascular barriers that limit traditional nanomedicines, offering a direct, non-invasive route to the primary tumor site [140], [141]. Once the AuNP-carrying probiotics colonize the colorectal lesions, localized near-infrared (NIR) irradiation can be applied to trigger highly specific photothermal ablation, sparing the surrounding healthy intestinal mucosa [142].

However, transforming wild-type or standard probiotics into highly efficient nanocarriers presents significant bio-engineering hurdles. A critical challenge lies in the immobilization of AuNPs onto or within the bacterial chassis without compromising cellular viability, motility, or tumor-homing fidelity. Current research explores two main methodologies to achieve this integration:

#### Surface Bio-conjugation

Utilizing electrostatic interactions, covalent coupling, or specific biotin-streptavidin bonds to attach pre-synthesized AuNPs to the bacterial outer membrane or flagella [143].

#### Intracellular Biosynthesis

Engineering or exploiting inherent metabolic pathways to allow the bacteria to reduce gold salts and synthesize AuNPs *in situ* intracellularly [144].

Optimizing the cargo-loading capacity while ensuring the biosecurity, controlled clearance, and predictable *in vivo* behavior of these living materials remains the frontier challenge in realizing the full potential of probiotic-mediated photothermal therapies.

In this research, we leverage EcN as a living host chassis to synthesize, accommodate, and deliver gold nanoparticles, addressing the critical balance between nanoparticle yield and carrier biomotility.

The choice between extracellular and intracellular routes fundamentally dictates the downstream processing and physicochemical characteristics of the colloids. Table 4.1 summarizes the advantages and limitations of both routes.

**Table 4.1.** Advantages and challenges of extracellular versus intracellular microbial biosynthesis pathways.

| **Biosynthesis Route** | **Mechanism** | **Key Advantages** | **Major Challenges** |
|---|---|---|---|
| **Extracellular** | Exploits secreted enzymes, proteins, and organic metabolites to reduce metal ions in the surrounding aqueous medium. | Simplifies downstream recovery and purification of nanoparticles without cell lysis. | Often results in wider size distributions and lower control over nanoparticle morphology. |
| **Intracellular** | Involves the active or passive uptake of metallic ions Au<sup>3+</sup> across the cell envelope into the cytoplasm, followed by enzymatic reduction. | Produces highly monodisperse, crystalline particles at a lower nanometer scale (typically 10-20nm). | Requires complex extraction protocols if nanoparticle isolation is needed, increasing processing costs. |

### Green Synthesis and Intracellular vs. Extracellular Biosynthesis of AuNPs in EcN for Enhanced PTT

**T**o overcome the biocompatibility issues and toxic chemical residues associated with conventional nanoparticle synthesis, this study explores an eco-friendly and biosafe strategy by developing a bio-hybrid platform utilizing EcN integrated with AuNPs obtained via green chemistry. Typically, chemical methods employ hazardous reducing and stabilizing substances like sodium borohydride or cetyltrimethylammonium bromide (CTAB). To ensure that the resulting AuNPs maintain negligible cytotoxicity and pristine surface properties highly compatible with living bacterial systems, we utilize a facile, room-temperature green synthesis approach. This method utilizes gold chloride (HAuCl_4_) as the metallic precursor and ascorbic acid (Vitamin C) as a sustainable, non-toxic reducing and stabilizing agent [145]. By optimizing the stoichiometric ratio of gold chloride to ascorbic acid, AuNPs can be fabricated through microbial routes, which are broadly categorized into extracellular and intracellular pathways.

### Mechanisms and Dynamics of Intracellular AuNPs Formation

The intracellular synthesis of gold nanoparticles (AuNPs) within microbial hosts proceeds through a well-defined three-step biochemical mechanism: i) the active or passive transport of Au^3+^ ions into the cytoplasm via specific membrane transporters or localized surface adsorption; ii) intracellular reduction mediated by NADH-dependent reductases or small-molecule endogenous reductants such as reduced glutathione (GSH); and iii) intracellular nucleation and subsequent crystal growth within the confined cellular microenvironment [146], [147], [148]. This metabolic process often yields nanoparticles that are tightly associated with cellular components, such as periplasmic proteins and inner membrane lipids. This intimate association provides a natural, robust biomolecular capping layer that confers exceptional colloidal stability, prevents irreversible aggregation, and ensures high biocompatibility for downstream biomedical applications.

However, executing this intricate biosynthetic process to engineer functional, living “bacteriobots” requires maintaining a delicate homeostatic equilibrium. High intracellular concentrations of gold ions can exert severe cytotoxic effects by inducing oxidative stress, disrupting the transmembrane potential, and compromising bacterial viability and flagellar motility. Therefore, critical physicochemical parameters, including precursor metal ion concentration, incubation exposure time, bacterial growth phase, pH, and ambient temperature, must be precisely optimized to maximize nanoparticle yield while strictly preserving the biological and structural integrity of the living carrier.

In this context, the selection of *Escherichia coli* Nissle 1917 (EcN) as the biological factory is strategically dictated by its unique physiological profile as a safe, non-pathogenic probio-bot. Beyond the implementation of this living platform, a truly sustainable nanobiotechnology paradigm dictates the elimination of harsh synthetic chemicals (such as sodium borohydride or hydrazine) by introducing eco-friendly, “green” auxiliary substances. In this study, the integration of ascorbic acid as a biocompatible, green co-reductant represents a crucial milestone in optimizing the biosynthetic route.

The deployment of ascorbic acid within this bacterial matrix serves a dual chemical and structural purpose, operating as a controlled, secondary reducing agent. While the endogenous metabolic machinery of living EcN successfully drives the initial internalization and core nucleation (Au^3+^ → Au^0^), the intrinsic intracellular reduction capacity of the bacteria may face thermodynamic limitations when clearing high concentrations of residual precursor gold ions. The addition of ascorbic acid, a mild and non-toxic antioxidant, supplements this process by providing an eco-friendly reservoir of electrons. Because ascorbic acid acts as a weak reducing agent under ambient conditions, it lacks the chemical potential to trigger spontaneous, chaotic extracellular nucleation in fluid phases lacking pre-existing crystal faces. Instead, it selectively drives a seed-mediated growth mechanism, directing the reduction of remaining gold ions exclusively onto the intra-bacterial metallic seeds previously established by EcN.

Furthermore, this green substance plays a pivotal role in the optical and structural maturation of the nanostructures. The interaction between the intracellular gold nuclei, the surrounding bacterial biomolecules, and the ascorbic acid coordinates a synchronized crystallization. This geometric stabilization shifts and consolidates the characteristic Surface Plasmon Resonance (SPR) absorption band toward the optimal spectroscopic region (∼560 nm). By avoiding conventional toxic surfactants, this hybrid EcN-ascorbic acid tandem ensures that the entire life cycle of the nano-radiator remains strictly bio-friendly. The native bacterial envelope retains its structural integrity, acting as an organized micellar shield that provides a uniform protein coating (*capping*), which ultimately suppresses colloid aggregation and preserves the photothermal conversion efficiency required for effective photothermal therapy (PTT) against malignant cells.

## Materials and Methods

### Bacterial Strain and Culture Conditions

The probiotic model chasis, *Escherichia coli* Nissle 1917 (EcN) (Ardeypharm GmbH, Germany), was maintained as glycerol stocks at -80 °C. Prior to use, the bacteria were streaked onto Luria-Bertani (LB) agar plates (10 g/L tryptone, 5 g/L yeast extract, 10 g/L NaCl, and 15 g/L agar) and incubated at 37 °C for 24 h to obtain single colonies.

For liquid cultures, a single well-isolated colony of EcN was inoculated into 5 mL of sterile LB broth in a standard 15 mL conical tube. This preculture was incubated overnight (approx. 16 h) at 37 °C under continuous orbital shaking at 200 rpm to reach the stationary phase.

For experimental scaling or nanoparticle integration, the overnight culture was subcultured by diluting the bacterial suspension 1:100 into fresh, sterile LB medium in an Erlenmeyer flask (maintaining a 1:5 liquid-to-air volume ratio to guarantee optimal oxygenation). The culture was grown under the same environmental conditions (37 °C, 200 rpm) until reaching the stationary growth phase. Bacterial concentration was subsequently determined and standardized via colony-forming unit (CFU) counting on LB agar plates.

### Microbial Intracellular Synthesis of AuNPs Using EcN

To achieve the intracellular biosynthesis of gold nanoparticles (AuNPs) within the probiotic chassis, a sequential reduction protocol mediated by ascorbic acid was developed. Briefly, a 5 mL culture of *E. coli* Nissle 1917 (EcN) was prepared in sterile LB medium and incubated overnight at 37 °C under orbital shaking (200 rpm) to reach the stationary phase.

The bacterial suspension was centrifuged at 14,000rpm for 10 min to pellet the cellular biomass. To completely eliminate residual supernatant, metabolic byproducts, and extracellular materials that could interfere with the synthesis, the pellet was washed three times with 5 mL of sterile phosphate-buffered saline (PBS, pH 7.4). The resulting washed bacterial cells were resuspended in 5 mL of a 1 mM chloroauric acid (HAuCl_4_) aqueous solution (metallic precursor) and incubated overnight at room temperature (22 °C) under gentle agitation to allow the active diffusion and absorption of gold ions into the bacterial cells.

Following the 1-day precursor exposure, the cell suspension was centrifuged (14,000 rpm, 10 min) and washed three times with 5 mL of sterile PBS to thoroughly remove any unbound or extracellular gold salt ions. Next, the bacterial cells containing the intracellularly absorbed gold precursor were resuspended in 5 mL of an aqueous ascorbic acid (Vitamin C) solution, acting as a biosafe, non-toxic reducing and stabilizing agent) and incubated overnight at room temperature.

After 24 h of reduction, the bio-hybrid platform was subjected to a final purification cycle, consisting of centrifugation and three successive washes with 5 mL of sterile PBS to remove excess ascorbic acid and any loosely bound surface residues. The successful intracellular synthesis of gold nanoparticles within the bacterial vehicle (designated as EcN::AuNPs) was macroscopically monitored and confirmed by a distinct, irreversible color transition of the bacterial pellet and medium from pale yellow to a deep dark-brownish hue, a characteristic indicator of the localized surface plasmon resonance (LSPR) of mature Au^0^ nanoparticles.

## Characterization

### UV-Vis spectroscopy

The optical properties of synthesized bacterial gold nanoparticles in EcN were confirmed using UV-Vis spectrophotometer to detect the surface plasmon resonance (SPR) peak, confirming the formation of nanoparticles using a multi-modal reader BioTek Synergy HTX

### In Vitro Photothermal Therapy (PTT)

To evaluate the photothermal ablation efficiency of the engineered bio-hybrid platform, *in vitro* PTT assays were performed utilizing a localized near-infrared (NIR) laser system. The photothermal irradiation was systematically executed using the following optimized, fixed instrumental parameters: a beam spot area of 0.5cm^2^, an operating current of 2.8A, an output laser power of 0.915W, and a calculated power density (irradiance) of 1.83 W·cm^2^. The total continuous irradiation time was set to 5 min per treated well.

The experimental setup was conducted in multi-well plates, where cells EcN::AuNPs were exposed to the NIR laser. To ensure statistical validity and isolate the mechanisms of cellular death, the following cohort of experimental control groups was simultaneously evaluated under identical environmental conditions:

1. **Negative Control:** Untreated cells maintained under basal culture conditions (no nanoparticles, no laser exposure).
2. **Laser-Only Control:** Cells exposed to the NIR laser irradiation (1.83 W·cm^2^ for 5min) in the absence of any nanoparticle formulation, ruling out intrinsic laser-induced hyperthermia.

## Results

### Visual Confirmation and Optoelectronic Dynamics of AuNP Synthesis

At the experimental level, the preliminary confirmation of successful gold nanoparticle (AuNP) formation within the *Escherichia coli* Nissle 1917 (EcN) matrix was established through macroscopical visual observation. Upon consecutive exposure of the washed bacterial biomass to the HAuCl_4_ precursor and the green reducing agent ascorbic acid, a highly distinct and rapid chromatic shift was observed. Figure 1 shows the color change associated with AuNP biosynthesis. This macroscopical color change serves as a reliable, direct diagnostic indicator of the successful reduction of trivalent gold ions (Au^3+^) into elemental, metallic zero-valent gold nanoparticles (Au^0^) confined within the microbial hosts.

From a physical perspective, this fundamental optical phenomenon is governed by the Surface Plasmon Resonance (SPR) effect, a unique optoelectronic property that emerges exclusively when noble metal particles are downscaled to the nanometer range[149]. SPR occurs when the frequency of the incident electromagnetic fields (visible light) matches the collective, coherent oscillation of the free conduction electrons localized on the surface of the gold nanostructures, resulting in the selective absorption and scattering of specific wavelengths.

Because this synthesis takes place strictly via an intracellular pathway, the chromatic signature is deeply influenced by the biological environment. Once the Au^3+^ ions are actively or passively internalized, the abundant pool of cytosolic NADH-dependent reductases, chaperones, and small antioxidant molecules (such as reduced glutathione) initiate core nucleation. Subsequently, the addition of ascorbic acid as a green co-reductant drives a seed-mediated growth mechanism over those intracellular spots. As these crystals mature, the native proteome and phospholipid membranes of EcN immediately engulf the expanding metallic spheres. This continuous biomolecular coating acts as a natural *capping agent*, which provides a strong steric and electrostatic hindrance that effectively immobilizes the particles inside the cellular bounds, preventing fluid-phase aggregation and maintaining a stable, uniform size distribution. Consequently, this structural stability is manifested as a sharp, intense absorption band centered in the green-to-yellow spectrum (∼560 nm), transforming the living probio-bots into highly efficient nano-radiators tailored for targeted oncological hyperthermia.

**Figure 3.1.**
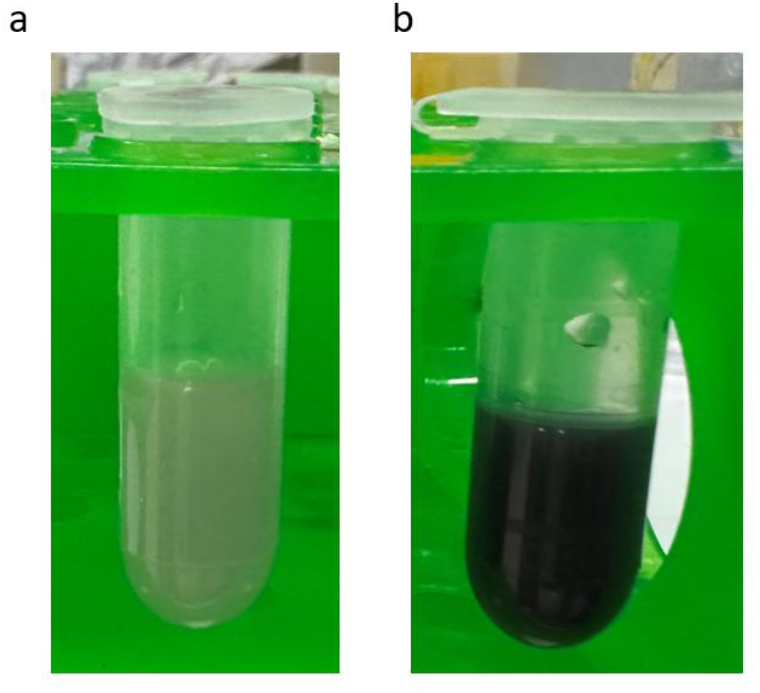
Color change associated with the biosynthesis of AuNPs. **(a)** E. coli Nissle 1917 culture. **(b)** E. coli Nissle 1917 culture after the addition of HAuCl₄ and ascorbic acid.

### UV-Vis Spectroscopic Characterization and Plasmonic Analysis

To rigorously confirm the intracellular reduction of trivalent gold into zero-valent crystalline nano-frameworks, UV-Visible spectrophotometric analyses were systematically performed. This technique serves as a fundamental, non-destructive diagnostic tool in nanobiotechnology due to its extreme sensitivity to the collective electronic state of noble metal boundaries. Figure 3.2 shows the different UV–Vis spectra measured.

**Figure 3.2.**
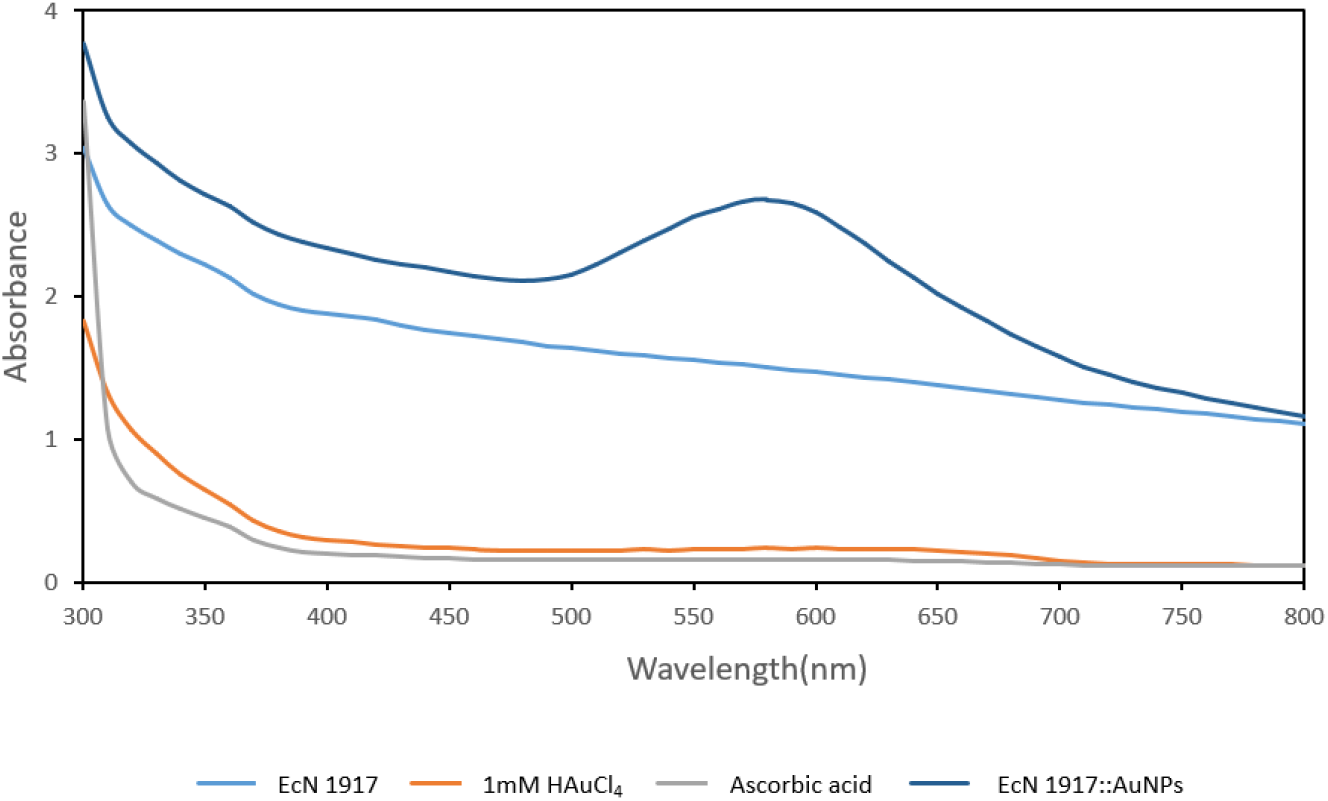
UV-Vis spectra.

The experimental spectra revealed a highly distinctive, intense, and prominent absorption band localized within the green-to-red visible region (spanning from 500 nm to 700 nm), exhibiting a well-defined maximum peak centered at 560 nm for the bio-hybrid system composed of *Escherichia coli* Nissle 1917 (EcN) treated with 1 mM HAuCl_4_ and supplemented with ascorbic acid.

Conversely, the negative control samples, consisting of pure, untreated EcN 1917 bacterial suspensions, aqueous solutions of unreduced 1 mM HAuCl_4_ precursor alone, and standalone ascorbic acid solutions, did not exhibit any discernible absorption peaks or plasmonic features within the monitored spectral window. The flat baseline associated with the controls confirms that neither the bare microbial biomass nor the molecular precursors possess intrinsic optical transitions in this region, proving unequivocally that the emergent peak at 560 nm is the direct and exclusive signature of the localized SPR of newly generated AuNPs.

The physical and structural implications of the observed SPR profile provide critical insights into the morphology and confinement environment of the biogenic AuNPs. The positioning of the resonance maximum at 560 nm is inherently linked to the size and shape of the synthesized structures. According to Mie’s scattering theory for spherical metallic particles, an SPR peak localized around 530–560 nm corresponds to stabilized, isotropic (spherical or quasi-spherical) gold nanostructures whose average core diameters fall within the approximately 30 to 60 nm range [149], [150], [151].

The fact that the experimental peak manifests as a noticeably broad band extending from 500 nm to 700 nm yields valuable thermodynamic and structural information. In classical colloidal chemistry, a highly symmetrical, ultra-narrow absorbance peak indicates a perfectly monodisperse population of isolated particles suspended in an ideal liquid continuum. Within our living probiotic platform, the broadening of the SPR band is a direct spectroscopic consequence of two concurrent factors:

1. **Intracellular Polydispersity:** The intracellular synthesis takes place within complex cellular compartments (periplasm and cytosol) where the distribution of endogenous reducing enzymes and cellular volumes varies slightly between individual cells, naturally leading to a controlled, moderate polydispersity in nanoparticle diameters.
2. **Biological Confinement and Aggregation Dynamics:** Because these AuNPs are tightly packed and confined within the highly crowded macromolecular matrix of the EcN microenvironment, the nanoparticles experience near-field electromagnetic coupling. When noble metal spheres are immobilized close to one another inside a cellular envelope, their individual plasmon fields interact, causing a characteristic broadening and a slight red-shift of the absorbance tail toward longer wavelengths [151].

From a therapeutic and translation perspective, this specific plasmonic profile is highly advantageous for oncological application. The wide absorption footprint covering a significant portion of the red and near-infrared (NIR) boundary ensures that the engineered EcN bacteriobots possess a high cross-section for light-to-heat conversion. When exposed to external laser sources, the broad plasmonic band guarantees efficient photon capture, facilitating rapid relaxation pathways that transfer energy into the surrounding tumor tissue. This validates the hybrid EcN-mediated green synthesis as an optimized platform for targeted photothermal ablation.

### Impact of Intracellular Synthesis on Cell Viability

A critical parameters for the real-world biomedical translation of engineered “bacteriobots” is the preservation of their biological viability. In our study, while the intracellular biogenic protocol achieved highly efficient reduction and stabilization of AuNPs (as confirmed by visual chromatic shifts and robust SPR profiles), a profound drop in *Escherichia coli* Nissle 1917 (EcN) survival was quantified. Specifically, the bacterial concentration collapsed from an initial post-culture density of 10^8^ CFU/mL in Luria-Bertani (LB) broth down to 10^3^ CFU/mL immediately following the completion of the synthesis protocol. This dramatic decline indicates that while EcN possesses the biochemical machinery required to act as an autonomous nanofactory, the localized accumulation of gold nanoparticles exerts severe, irreversible cytotoxic stress on the biological carrier.

From a mechanistic perspective, this massive loss of viability can be attributed to several synergistic factors operating during the intracellular nucleation and growth phases:

1. **Intracellular Disruption by Heavy Metal Toxicity:** The initial internalization of Au^3+^ ions from the HAuCl_4_ precursor triggers immediate heavy metal shock. Trivalent gold is a powerful oxidizing agent that binds non-selectively to sulfhydryl (-SH) groups in cytosolic proteins, enzymes, and structural components of the bacterial cell envelope. This interaction inactivates essential metabolic pathways, cross-links vital enzymes, and compromises membrane potential before complete reduction to Au^0^ can take place[152].
2. **Mechanical Disruption and Spatial Confinement:** As the nanoparticles mature and grow inside the restricted volumes of the periplasm and cytoplasm, they exert mechanical pressure against the inner membrane and the peptidoglycan cell wall. The physical crowding of hundreds of metallic nano-spheres within a single micro-organism alters structural integrity, leading to micro-ruptures in the cellular envelope and triggering autolysis in the majority of the population.
3. **Severe Metabolic and Oxidative Stress:** The massive mobilization of endogenous reducing agents, such as NADH-dependent reductases and reduced glutathione (GSH), to reduce Au^3+^ rapidly exhausts the cell’s metabolic currency. This sudden depletion of the pool of cellular reducing equivalents deprives the bacteria of the ability to manage reactive oxygen species (ROS) or maintain ion homeostasis, driving the cells into a terminal energetic crisis.

Despite this high mortality rate, the remaining population of surviving cells 10^3^ CFU/mL holds significant therapeutic value. Literature in advanced nanomedicine demonstrates that for targeted cancer applications, a dense, overpopulated bacterial vector can sometimes trigger an excessive and dangerous systemic immune response (e.g., cytokine storms) *in vivo*. Therefore, a bio-hybrid formulation containing highly stable, intra-bacterially confined AuNPs carried by a strictly regulated, lower concentration of living, motile probio-bots represents an optimized equilibrium. The 10^3^ CFU/mL cohort retains the necessary active tumor-homing and chemotactic capabilities toward the hypoxic cores of solid tumors, while the protective “capping” matrix provided by the lysed bacterial biomass prevents the systemic aggregation of the released nanoparticles, maintaining their long-term colloidal stability for subsequent photothermal ablation.

### Evaluation of Photothermal Therapy (PTT) and Laser-Induced Hyperthermia Kinetics

To validate the translational efficacy of the engineered *Escherichia coli* Nissle 1917 bio-hybrids loaded with intracellular gold nanoparticles (EcN::AuNPs), their photothermal conversion capacity was systematically evaluated under near-infrared (NIR) laser irradiation. The application of light within the NIR window is a core requirement for non-invasive oncological therapies, as it minimizes photon scattering and competitive absorption by endogenous tissue chromophores such as hemoglobin and water, allowing deeper penetration into biological matrices.

The experimental PTT setup was conducted using a continuous-wave NIR laser system operating at an electrical current of 2.8 A, calibrated to deliver an optical output power of 0.915 W. Focusing this beam onto a well surface area of 0.5 cm² yielded a precise and constant power density (irradiance) of 1.83 W/cm². This specific energy density was maintained throughout an exposure window of 5 minutes (300 seconds) per sample.

Upon continuous laser exposure, both the standalone biosynthesized AuNPs and the EcN::AuNPs-treated formulations demonstrated a significant, concentration-dependent escalation in temperature. Conversely, the negative control samples (untreated cells or vehicle blanks) exhibited negligible thermal shifts under identical irradiation conditions. This stark contrast confirms that the macroscopical heating is driven exclusively by the localized Surface Plasmon Resonance (SPR) of the internal gold nanostructures, which efficiently absorb the incident photons and rapidly dissipate that energy into the immediate fluid continuum via non-radiative relaxation pathways.

### Steady-State Temperature Shifts and Real-Time Hyperthermia Kinetics

The thermal profiles of the intracellular biogenic AuNPs across varying concentrations were characterized using two distinct analytical viewpoints to fully capture their nano-radiator performance.

Figure 3.3 outlines the absolute temperature shift of EcN::AuNPs from the baseline value to the final equilibrium state captured exactly at the 5-minute endpoint. At the highest tested concentration, the applied power density of 1.83 W/cm² was sufficient to induce a drastic thermal escalation, driving the formulation from a physiological baseline temperature of 20°C before laser irradiation up to a maximum peak of 75.4°C after 300 seconds. This total temperature increment (ΔT≈ 55.4°C) is remarkably high and vastly exceeds the classical hyperthermia threshold (42–47°C), successfully entering the regime of rapid thermal ablation capable of causing immediate physical damage, protein denaturation, and localized destruction of surrounding malignant structures. As controls, EcN culture alone, HAuCl₄ alone, and ascorbic acid alone were evaluated. No significant ΔT was observed in any of these conditions.

**Figure 3.3.**
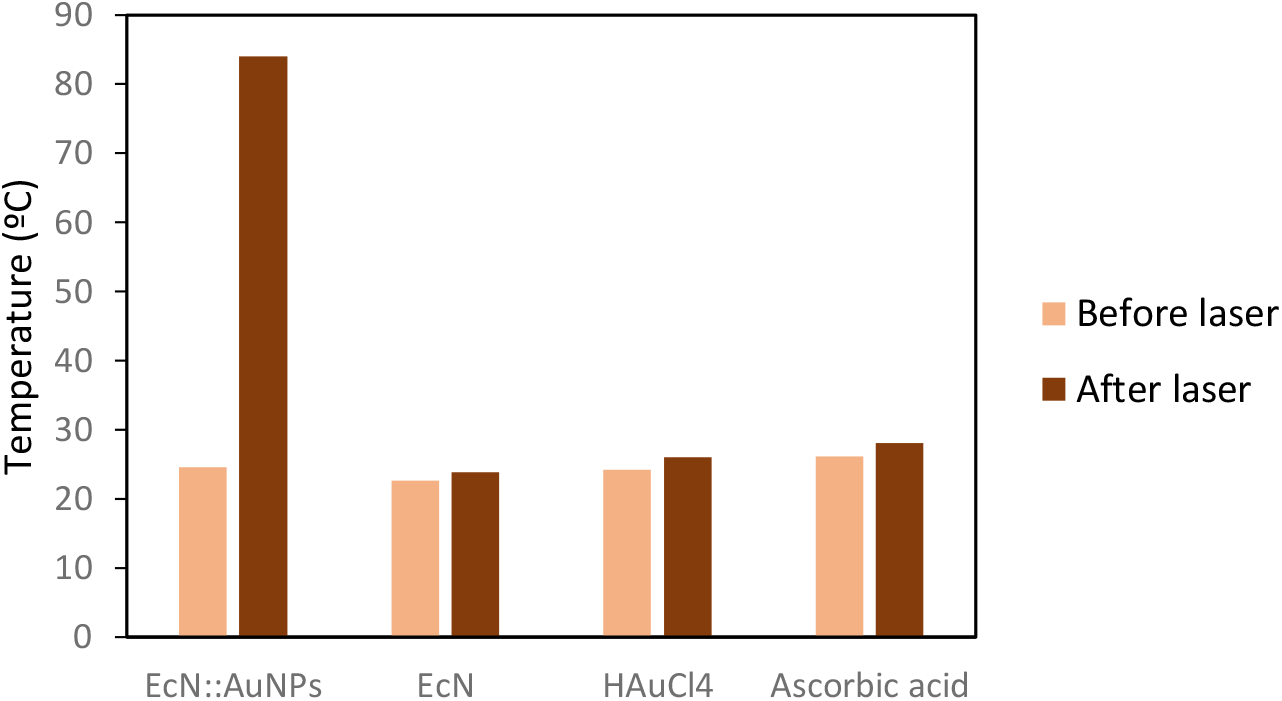
Kinetic temperature profiles of different samples, including the EcN::AuNPs, EcN alone, the HAuCl_4_ precursor alone, and the ascorbic acid reducing agent alone, before and after near-infrared (NIR) laser irradiation for 300 seconds.

### Real-Time Kinetic Profiling and Thermal Lag Phase Phenomena

To elucidate the velocity and thermodynamic behavior of this light-to-heat conversion, the continuous temperature shifts of the various EcN::AuNPs formulations were tracked, recording definitive data points after each 30-second interval across the 300-second treatment window. In this kinetic profile, a concentration value of 100% represents the stock, undiluted EcN biological culture immediately after completing the intracellular gold nanoparticle synthesis protocol. The subsequent percentages correspond to systematic serial dilutions of this original stock culture in ultra-pure Milli-Q water, while the 0% control represents a bare EcN bacterial culture subjected to identical washing steps but lacking any internal AuNPs.

The resulting real-time kinetic curves reveal two distinctly different thermodynamic regimes depending on the concentration of the nano-radiators:

**Figure 3.4.**
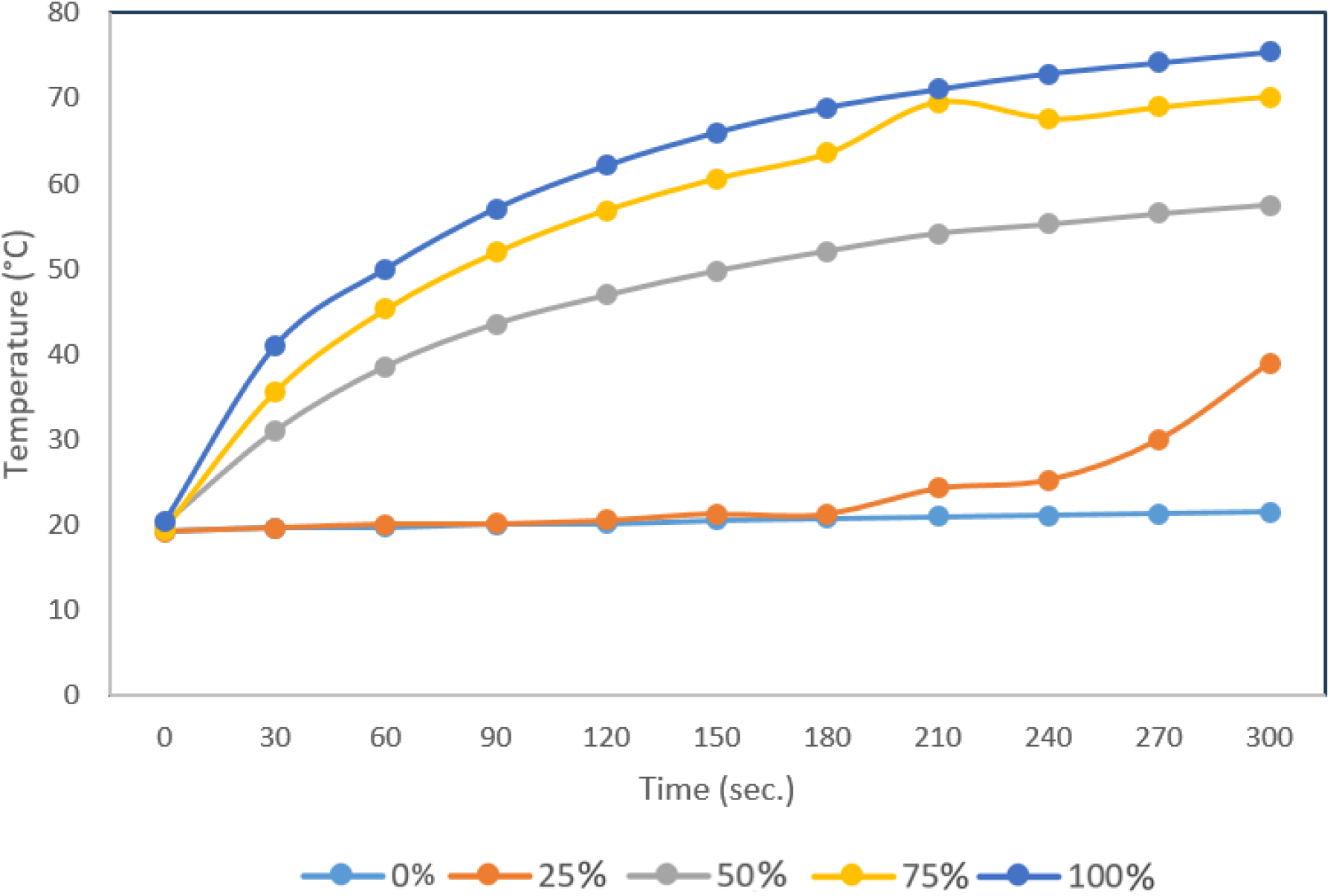
Real-time kinetic profiling of the photothermal temperature increase induced by different concentrations of EcN::AuNPs.

#### 1. High-Concentration Regime (relative concentracion>50%)

At elevated concentrations (>50%), the *EcN::AuNPs* system exhibits an immediate, progressive, and highly linear temperature increase from the very onset of laser exposure (t = 0). This instantaneous thermal response indicates that the density of intracellular AuNPs is high enough to immediately overcome the specific heat capacity of the surrounding aqueous medium. The high volumetric concentration of plasmonic cores guarantees a massive, simultaneous cross-section of photon absorption, driving a rapid and continuous non-radiative relaxation that manifests as an immediate macroeconomic temperature rise.

#### 2. Low-Concentration Regime and the Thermal Lag Phase (<50%)

In contrast, at lower operational concentrations, the system displays a peculiar non-linear kinetic profile characterized by a prolonged initial period with no detectable temperature increase, followed by a delayed, progressive thermal escalation later in the treatment. This distinct “lag phase” at low nanoparticle densities can be scientifically interpreted through a combination of threshold thermal mass and local-to-global heat diffusion dynamics:

- **The Heat Sink Effect of the Solvent Matrix:** Water possesses an exceptionally high specific heat capacity (4.184 J·Kg^-1^·K^-1^). At high dilutions, the absolute number of EcN::AuNPs per unit volume is low, meaning the total localized thermal energy generated during the first few seconds of laser irradiation is entirely consumed by the immediate surrounding water molecules acting as a thermal “sink.” Until the collective micro-environmental temperature of these hydration shells reaches a specific thermodynamic threshold, no macroscopical temperature change can be registered by the measurement system.
- **Near-Field vs. Far-Field Interparticle Distance**: At higher concentrations, the proximity between individual EcN cells packed with AuNPs is minimal, enabling rapid near-field plasmonic and thermal coupling where overlapping heat zones cause an exponential cumulative rise. At low concentrations, however, the distance between the nano-radiators increases significantly. Heat dissipation from an isolated nanostructure drops sharply as a function of distance (1/r). Consequently, a prolonged period of continuous laser pumping is mandatory for these scattered thermal fronts to diffuse through the cold solvent gaps, collide, and finally trigger a measurable global temperature increment.
- **Pericellular Thermal Envelopes**: It is highly probable that during the initial phase of irradiation at low concentrations, localized hyperthermia is actively occurring *within* and immediately around the EcN cell walls, yet this energy remains transiently confined at the microscale. Only after the continuous absorption of NIR photons induces a critical thermal breakdown of the protective bacterial membrane does this trapped kinetic energy burst into the bulk solution, causing the sudden, progressive macroscopical temperature rise observed in the latter half of the experiment.

This interpretation underscores that while lower concentrations of EcN::AuNPs require a longer laser-activation threshold to show global effects, they still retain significant photothermal conversion efficacy once this initial thermal inertia is successfully overcome, a factor that must be precisely calibrated when designing localized *in vivo* oncological protocols.

### Quantitative Evaluation of Photothermal Conversion Efficiency

To mathematically quantify the light-to-heat conversion performance of the engineered bio-hybrids, the photothermal conversion efficiency (η) of the EcN::AuNPs suspension under 806nm NIR laser irradiation (1W/cm^2^, beam area of 0.5cm^2^, net incident power $ = 0.50W) was calculated according to the equilibrium energy balance model developed by Roper et al. (2007)[153]. To account for the cylindrical optical geometry of the laser beam within the 1 mL sample vial, a localized thermodynamic boundary was defined utilizing an effective mass of 0.32g directly interacting within the irradiation pathway. First, the systemic heat transfer coefficient (hS) was determined during the passive cooling phase following laser deactivation. Given the recorded thermal decay time constant (τ_S_) of 310 sec. and the specific heat capacity of the aqueous matrix (C_p_ = 4.184 J·Kg^-1^·K^-1^), the heat dissipation parameter was calculated as hS = m_effective_ C_p_/τ_S_ = 0.004319W·C^-1^. Substituting this value into the steady-state balance equation, alongside the net temperature escalation (ΔT=55.4°C, from 20.0°C to 75.4°C), the measured optical absorbance (ABS_806_ = 0.523, yielding a fraction of absorbed light of 0.70), and the baseline solvent heat dissipation (Q_solvent_=0.010W), the photothermal conversion efficiency of the EcN::AuNPs was determined to be 65.5%.

This remarkably high efficiency significantly outperforms conventional chemically synthesized gold nanostructures or raw metallic formulations, which typically oscillate between 40% and 55% in the NIR window[154]. This enhanced performance can be attributed to the dense spatial confinement of the gold cores within the structural domains (periplasm and cytoplasm) of *E. coli* Nissle 1917. This intracellular crowding promotes constructive near-field plasmonic coupling[155], which maximizes the optical absorption cross-section and optimizes thermal transport to the surrounding environment without the macroscopical aggregation constraints that typically compromise raw colloidal systems in biological buffers.

